# Molecular-scale visualization of sarcomere contraction within native cardiomyocytes

**DOI:** 10.1101/2020.09.09.288977

**Authors:** Laura Burbaum, Jonathan Schneider, Sarah Scholze, Ralph T Böttcher, Wolfgang Baumeister, Petra Schwille, Jürgen M Plitzko, Marion Jasnin

## Abstract

Sarcomeres, the basic contractile units of striated muscle, produce the forces driving muscular contraction through cross-bridge interactions between actin-containing thin filaments and myosin II-based thick filaments. Until now, direct visualization of the molecular architecture underlying sarcomere contractility has remained elusive. Here, we use *in situ* cryo-electron tomography to unveil sarcomere contraction in frozen-hydrated neonatal rat cardiomyocytes. We show that the hexagonal lattice of the thick filaments is already established at the neonatal stage, with an excess of thin filaments outside the trigonal positions. Structural assessment of actin polarity by subtomogram averaging reveals that thin filaments in the fully activated state form overlapping arrays of opposite polarity in the center of the sarcomere. Our approach provides direct evidence for thin filament sliding during muscle contraction and may serve as a basis for structural understanding of thin filament activation and actomyosin interactions inside unperturbed cellular environments.

## Introduction

Muscle cells contain numerous myofibrils composed of adjoining micrometer-sized contractile units called sarcomeres (1). These large macromolecular assemblies feature thin filaments, made of polar actin filaments (F-actin) in complex with troponin (Tpn) and tropomyosin (Tpm), and bipolar thick filaments consisting of a myosin tail backbone decorated with myosin-II heads on either side of a head-free region (the so-called bare zone) (1). The sliding filament theory provided a framework for understanding the mechanism of muscle contraction based on the relative sliding between the myofilaments via cyclic interactions of myosin heads with F-actin (2–5). Five decades after these pioneering studies, much remains to be discovered at the structural level about how the molecular players of the actomyosin machinery co-operate within sarcomeres to produce contractile forces.

Muscle research has benefited greatly from the development of powerful X-ray sources and electron microscopes, allowing detailed views into muscle organization, time-resolved X-ray diffraction studies of contracting muscle, and three-dimensional (3D) reconstruction of actomyosin interactions (6–10). Thick filaments are known to be crosslinked in the central M-line and occupy the A-band (fig. S1A) (1, 11, 12). On either side of the M-line, interdigitating myofilaments organize into hexagonally packed arrays (1, 13, 14). Thin filaments extend throughout the I-bands toward the Z-disk, in which their barbed ends are anchored and crosslinked by α-actinin (fig. S1A) (1, 11, 12, 15, 16). At the molecular level, Ca^2+^-binding to Tpn is thought to trigger a shift in the azimuthal position of Tpm on F-actin to uncover the myosin binding site which allows actomyosin interactions (17, 18). Yet, none of the approaches used so far has permitted molecular-scale imaging of sarcomere organization inside unperturbed cellular environments. Recent advances in cryo-electron tomography (cryo-ET) provide access to the 3D molecular architecture of the cellular interior (19, 20). Here, we use state-of-the-art cryo-ET methodologies to unravel sarcomere organization in frozen-hydrated neonatal cardiomyocytes across length scales, from the myofilament packing to the sliding and functional states of the thin filaments enabling contraction.

## Results

Cardiomyocytes isolated from neonatal rat ventricles exhibit irregular, star-like morphology and spontaneous rhythmic contractions (movie S1). To explore the microscale organization of myofibrils mediating cell contractility, neonatal rat cardiac myocytes were immunolabelled for Tpn I (one of the three subunits of the Tpn complex), the heavy chain of cardiac myosin and α-actinin, as markers for the thin filament, the thick filament and the Z-disk, respectively. Myofibrils align into bundles along the principal axes of the neonatal cells (Fig. 1A and fig. S1, B and C). They feature regularly spaced Z-disks and A-bands, with a sarcomere length of 1.8 ± 0.2 μm (Fig. 1B and fig. S1D), similar to the value we found in adult mouse cells.

**Fig. 1.**
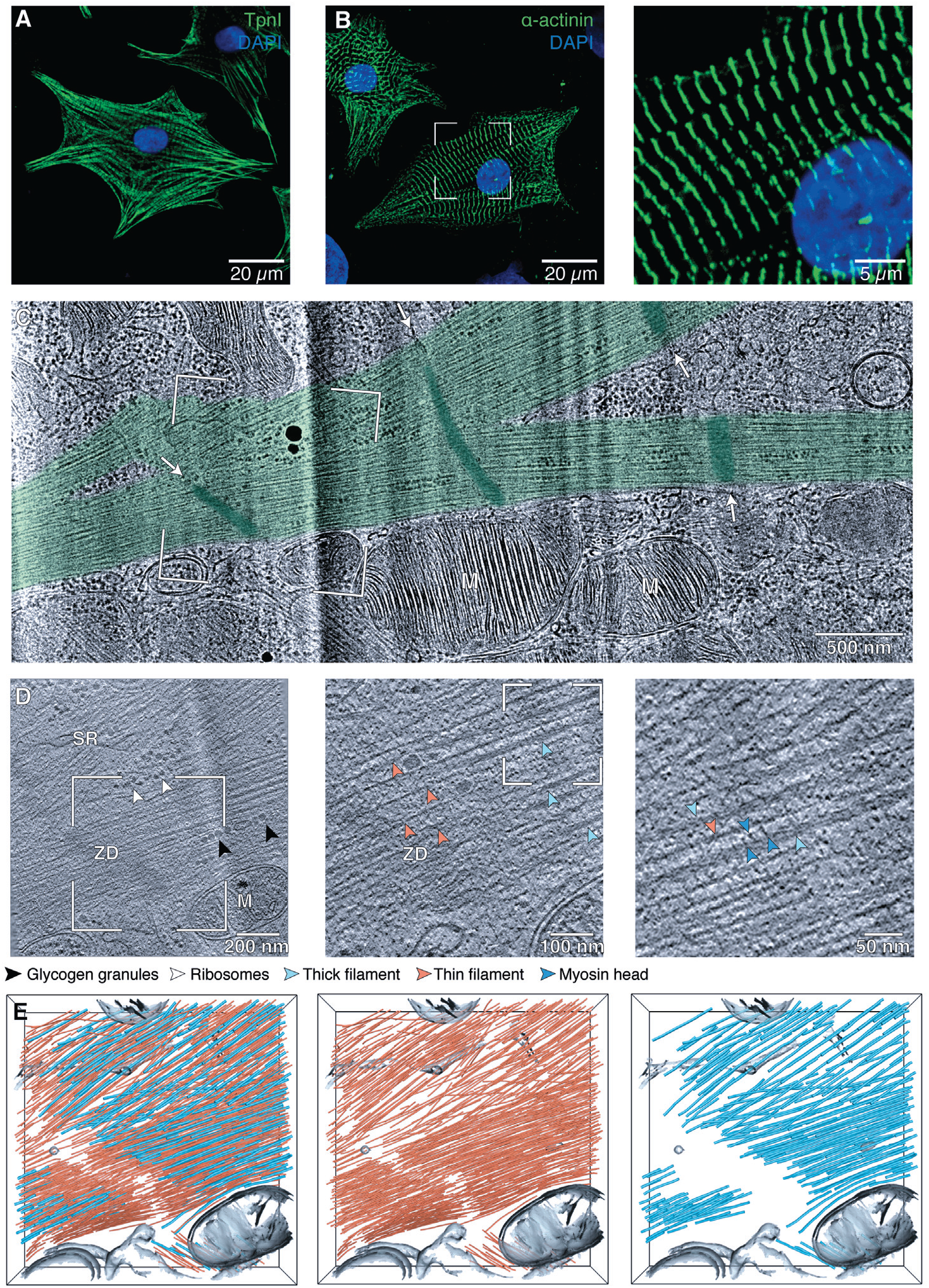
Visualizing the myofibrillar interior *in situ*. (A-B) Myofibrils (A) in neonatal rat cardiomyocytes align with the main axes of the star-like shaped cells and feature regularly spaced Z-disks (B). (See fig. S1, B-D for more examples.) (C) TEM image of the enlarged framed area of a neonatal rat cardiomyocyte lamella shown in fig. S2D. Myofibrils (green) display periodic, electron-dense bands (dark green, white arrows) merging across adjoining parts. M, mitochondria. (D) Slice from a tomographic volume acquired at the framed area in (C), showing the myofibrillar architecture. SR, sarcoplasmic reticulum; ZD, Z-disk. Left inset: Z-disk periphery. Right inset: myosin heads extending toward thin filaments. (E) 3D rendering of the cellular volume shown in (D) revealing the nanoscale organization of thin (orange) and thick (cyan) filaments within unperturbed sarcomeres. (See also movie S2). Additional examples are provided in fig. S3.

Unlike adult cardiomyocytes, the cell size is suitable for cryofixation by plunge-freezing (fig. S2, A and B). Neonatal cardiomyocytes were cultured on EM grids and plunge-frozen during spontaneous contraction (fig. S2B). 100-to 200-nm-thick vitrified cellular sections (so-called lamellas) were prepared using cryo-focused ion beam (cryo-FIB) milling (fig. S2C) (21). Transmission electron microscopy (TEM) images revealed cytoplasmic regions filled with long adjacent myofibrils surrounded by mitochondria and sarcoplasmic reticulum (Fig. 1C and fig. S2D). These myofibrils have a typical width of 470 ± 60 nm and display periodic, electron-dense bands that merge across adjoining parts (Fig. 1C and fig. S2D). The periodicity is comparable to the sarcomere length, suggesting that these bands may be either Z-disks or M-lines.

Myofibrils identified in the lamellas were imaged in 3D by cryo-ET. The reconstructed volumes permitted visualization of the thin and thick filaments, with diameters of 8.4 ± 0.9 nm and 17.4 ± 3.0 nm, respectively (Fig. 1D and fig. S3, A-C). Densities resembling the head domains of myosin II (10) bridge the space between interdigitating cardiac myofilaments (Fig. 1D, right). Macromolecular complexes of 34.4 ± 4.3 nm in diameter, most likely glycogen granules (22), are found outside and within the myofibrillar interior, whereas mitochondria, sarcoplasmic reticulum and ribosomes are found exclusively in the cytoplasmic space devoid of myofibrils (Fig. 1D and fig. S3, A-C).

The periodic, electrondense bands observed in the TEM images are made of irregular, fuzzy structures that intersect with the thin filaments (Fig. 1D and fig. S3, A-C). Filament segmentation provided a first glimpse at the native myofilament organization in 3D, revealing a gap between the thin filaments at the center of the band and the absence of thick filaments (Fig. 1E, fig. S3, D-F, and movie S2). Thus, the electron-dense structures correspond to Z-disks, with widths between 100 and 140 nm as has been reported for cardiac muscle (23, 24), and the regions depleted of thick filaments are I-bands, with lengths between 240 and 300 nm. Interestingly, we did not observe any myosin bare zone depleted of thin filaments (Fig. 1E, fig. S3, D-F, and movie S2).

We next quantitatively analyzed the 3D organization of the neonatal myofilaments using an approach described previously (25). Thick filaments organize in a hexagonal lattice with an interfilament distance of 45.1 ± 3.8 nm (Fig. 2A and fig. S4A). Thin filaments are found at the trigonal positions of the lattice, at a distance of 26.0 ± 2.4 nm from the thick filaments (Fig. 2B and fig. S4, B and C). However, the mean distance of 15.5 ± 1.4 nm between thin filaments indicates that they are also present outside the trigonal positions of the array (fig. S4D). This is in agreement with the ratio of thin-to-thick filaments of 3:1 found in these cells, as compared to the ratio of 2:1 reported in vertebrate muscle (26). Therefore, the packing of the thin filaments around the thick filaments we observed, both in real cross-sections and quantitatively, is less ordered than that found in skeletal muscle using X-ray diffraction (27–30) (Fig 2C and fig. S5). This indicates that cardiac sarcomere contraction is already possible with imperfect packing of the thin filaments, and suggests that thick filaments are an important determinant of myofilament organization.

**Fig. 2.**
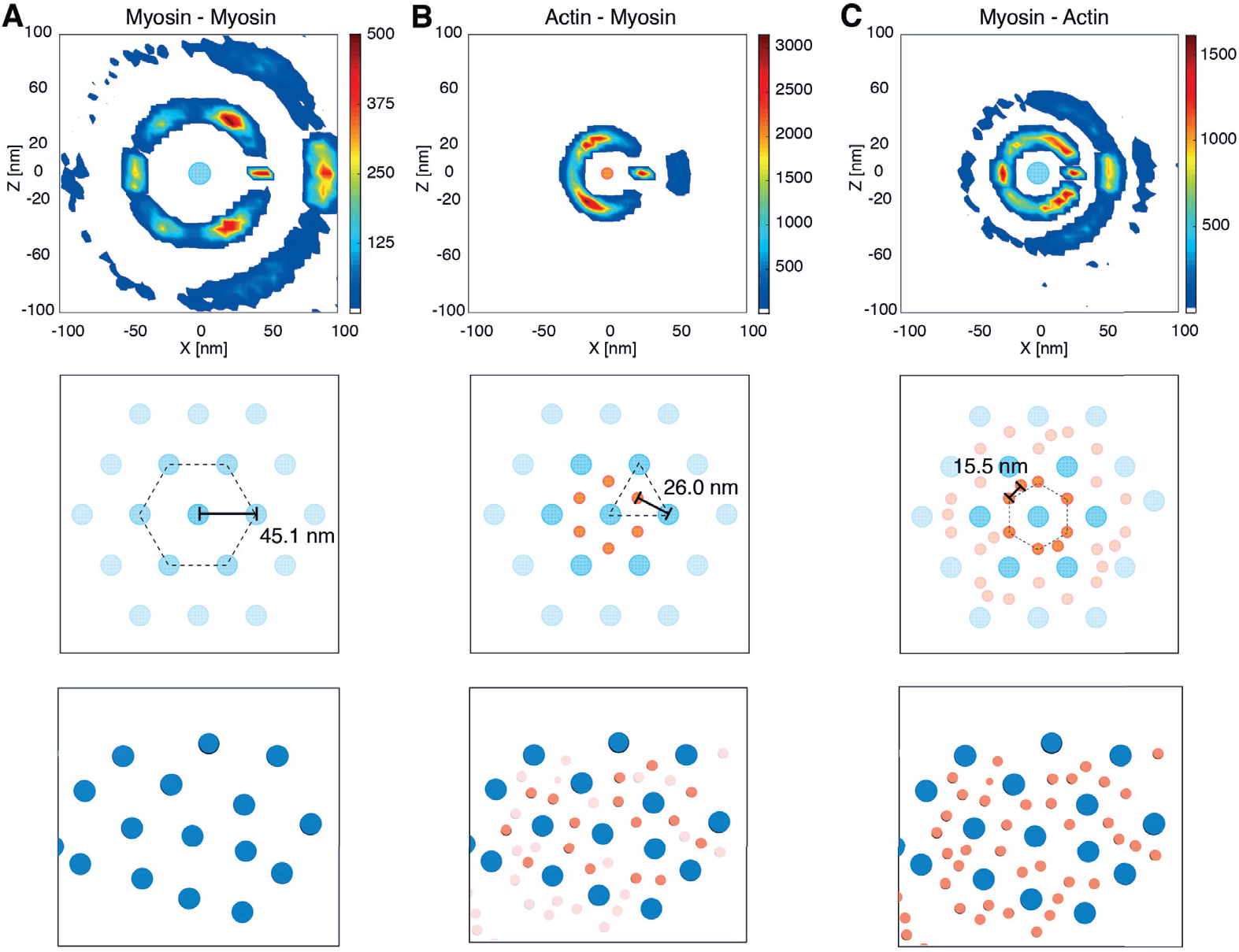
Neonatal cardiac thick filaments are hexagonally packed with an excess of thin filaments outside the trigonal positions. (A-C) Quantitative analysis of the myofilament packing in neonatal rat cardiomyocytes (top, see methods in SM), schematic representation of the array (middle), and cross-section through the myofibril shown in fig. S3E (bottom). See fig. S5 for additional cross-section examples. Heat map of nearest neighbor positions around thick filaments (“Myosin”, A and C, cyan-filled circle) or thin filaments (“Actin”, B, orange-filled circle) in a plane perpendicular to the filament cross-sections and within the distance ranges determined in fig. S4. Thick filaments assemble into a hexagonal lattice with an interfilament distance of about 45 nm (A and fig S4A). Thin filaments are found at about 26 nm from three neighboring thick filaments (B and fig S4, B and C) and outside the trigonal positions of the lattice (C and fig S4D).

Next, we exploited subtomogram averaging (31) to unveil the molecular organization of the thin filaments enabling sarcomere contraction. To this end, we first developed an approach to obtain the structure of the thin filaments by aligning and averaging multiple consecutive actin subunits along the filaments without imposing any helical symmetry (see methods in supplementary materials (SM)). Thin filaments located on one side of a Z-disk were used to generate a *de novo* structure of F-actin in complex with Tpm resolved at 20.7 Å (fig. S6, A-C). At this resolution, the positions of the actin subunits with respect to the two Tpm strands allow to discriminate between F-actin structures of opposite polarity (fig. S6D). Comparison with a pseudoatomic model of the F-actin-Tpm complex (10) permitted to allocate the barbed (+) and pointed (−) ends of the actin filament (Fig. 3A and fig. S6E).

**Fig. 3.**
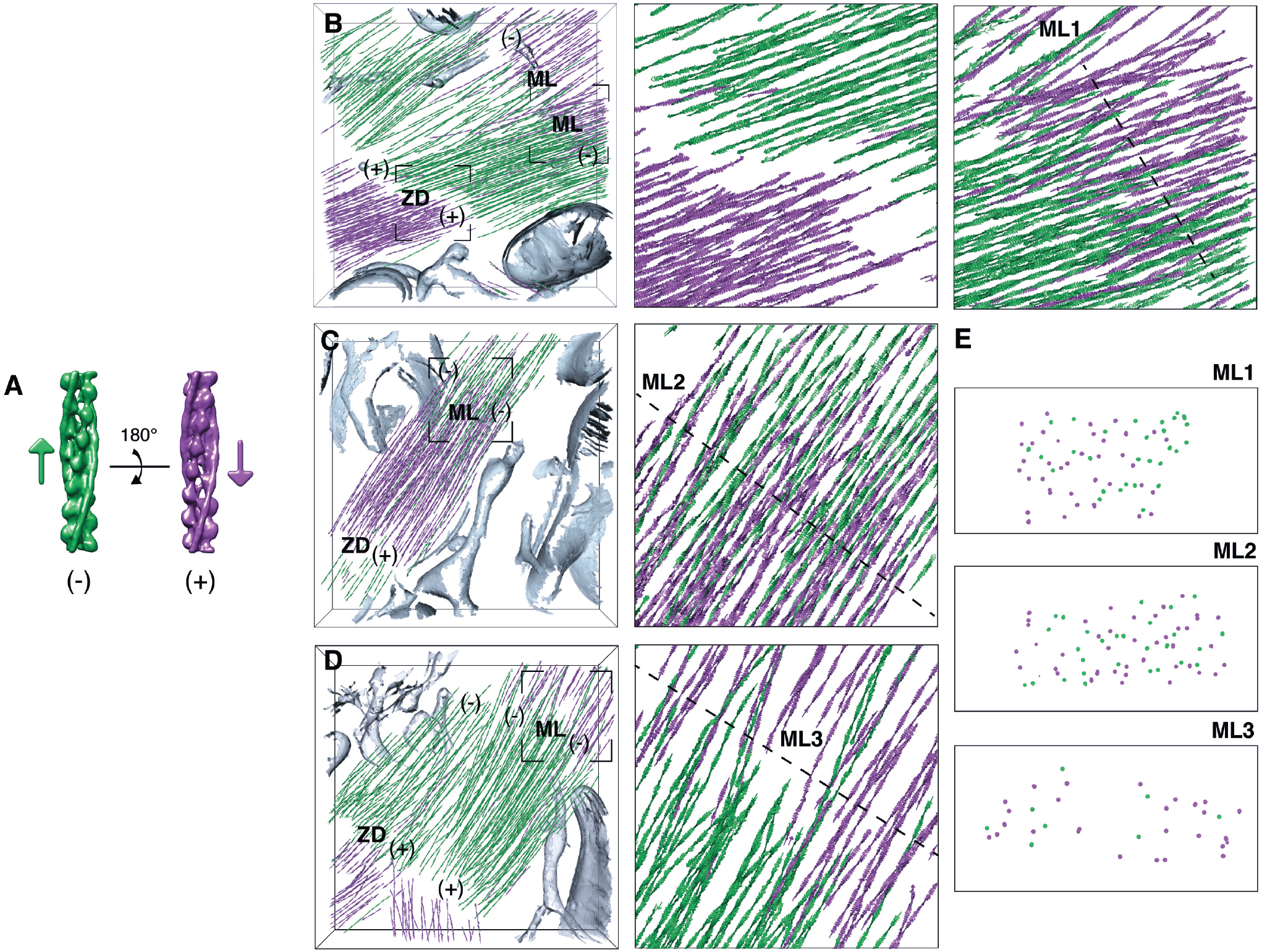
Structural signature of sarcomere contraction. (A) *De novo* structure of the actin-Tpm filament in opposite orientations, represented by arrows pointing toward the barbed (+) end. (B-E) Polarity assignment for the cardiac thin filaments shown in Fig 1E and movie S2 (B and movie S3) and fig. S3, E and F (C and D, respectively, and movie S4). Filaments are represented by arrows colored according to the assigned polarity. The gaps between thin filaments facing each other with their barbed ends indicate the location of Z-disks (ZD). The overlap regions between thin filaments of opposite polarity at the pointed (−) ends correspond to M-lines (ML). Insets: Thin filament orientations at the framed Z-disk (B, middle) and M-lines (B-D, right) with their respective cross-sections (E). See fig. S8 for visualization of the unassigned filaments.

We then sought to determine the polarity of the thin filaments within the native cardiac sarcomeres. Using a multireference alignment procedure based on the obtained *de novo* structure, we evaluated the polarity of each individual filament from a series of statistically independent estimates (see methods in SM; fig. S7, A and B). 69% of the thin filaments were assigned a polarity with confidence (fig. S7C) and visualized in 3D (Fig. 3). Filaments belonging to the same bundle shared the same polarity, while changes of polarity were observed between adjoining bundles (Fig. 3, B-D, left). In vicinity of the Z-disks, thin filaments face each other with their barbed ends, as has been previously reported (12, 15) (Fig. 3B, middle, and movie S3). Additionally, the analysis showed regions in which thin filaments of opposite polarity face each other with their pointed ends, revealing the location of M-lines (Fig. 3, B-D, left).

This allowed us to uncover the arrangement of the thin filaments in the center of the sarcomere, where arrays of opposite polarity are pulled toward each other during contraction. In two out of three M-lines, thin filaments of opposite polarity were found to overlap with each other, revealing their relative sliding (Fig. 3, B and C, and movie S4), as proposed by Gordon *et al*. (32). In the sarcomere shown in Fig. 3C, the overlap length is 281 nm, corresponding to 17% of the sarcomere length. This value is comparable to the values of 0.15 to 0.2 μm reported for the bare zone region (12), indicating that the sliding between these thin filaments occurs across the bare zone. The sarcomere length of 1.65 μm is smaller than the value of 1.8 ± 0.2 μm we measured in these cells, in agreement with sarcomere shortening. Cross-sections through the M-lines show that thin filaments of opposite polarities can contribute to the packing around the same thick filaments during their sliding (Fig. 3E, ML1 and ML2, and fig. S8, A and B). In the sarcomere shown in Fig. 3D, no clear overlap between thin filaments of opposite polarity was detectable at the M-line (Fig. 3, D and E, ML3, fig. S8C and movie S4), indicating that this sarcomere may be in a different contraction state.

We next used subtomogram averaging to determine the structures of the thin filaments associated to the observed M-line organizations. We first analyzed each tomogram individually in order to uncover potential differences in the Tpm position between sarcomeres. Tomograms acquired in cells where thin filaments of opposite polarity overlap in the M-line region provided a similar structure (Fig. 4E, left). In contrast, the data shown in Fig. 3D without clear overlap yielded a subtomogram average in which the Tpm density is shifted azimuthally on the surface of the actin filament (Fig. 4E), indicating that these filaments are in a different functional state.

**Fig. 4.**
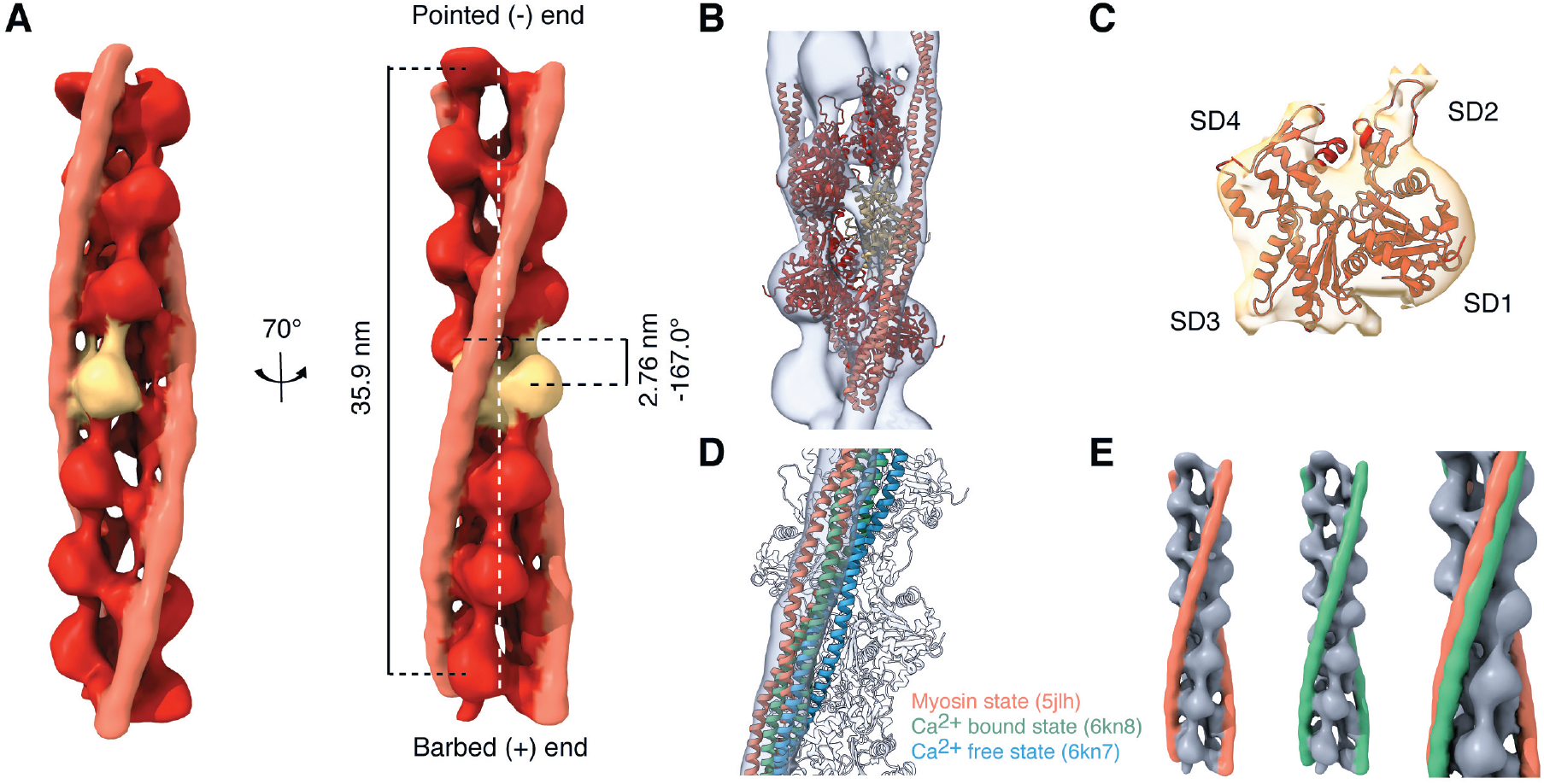
*In situ* structure of the cardiac thin filament in two distinct functional states. (A) *In situ* subtomogram average of the sarcomeric actin filament (red and yellow) in complex with Tpm (orange) resolved at 13.2 Å (fig. S9A, orange curve, movie S5). (B-C) Fit of a pseudoatomic model of the F-actin-Tpm complex (10) into the subtomogram average (B) reveals F-actin polarity at the level of the actin monomers (C). (D) Docking of pseudoatomic models in different states (10, 18) into the map reveals that the thin filament is in the myosin state (see also fig. S9E). (E) The thin filaments shown in Fig. 3D provided a structure in which the Tpm density (green) is shifted azimuthally as compared to the Tpm density obtained from the thin filaments shown in Fig. 3C (orange).

To generate a higher-resolution average, the tomograms with comparable structures were combined. A refined subtomogram average of the cardiac thin filament was obtained at a resolution of 13.2 Å (Fig. 4A, fig. S9, A and B, and movie S5). The structure features thirteen fully resolved F-actin subunits surrounded by two Tpm strands which approximately follow the same helical twist (Fig. 4A and movie S5). The actin filament consists of a left-handed single helix with a pitch of 35.9 nm, a rise per actin subunit of 2.76 nm and an angular twist per molecule of −167.0° (Fig. 4A, right), in agreement with the literature (33). The central part of the structure exhibits a local resolution of 11.4 Å (fig. S9C), allowing docking of a pseudoatomic model of the F-actin-Tpm complex into the map (Fig. 4B and movie S5). F-actin polarity is evident at the level of the actin monomers (Fig. 4C and movie S5). Comparison with pseudoatomic models in different functional states (10, 18) shows that the thin filament is in the fully activated (“myosin”) state that allows strong binding of the myosin heads (Fig. 4D and fig. S9, D, left, and E). This provides evidence at the molecular level that a majority of thin filaments in these sarcomeres are strongly bound to thick filaments, allowing their relative sliding and, thus, sarcomere contraction. Moreover, the Ca^2+^ bound state of the cardiac thin filament, which allows myosin head access but no strong binding (18), fitted best to the separate structure obtained from the data shown in Fig. 3D (fig. S9, D, right, and F), supporting that these sarcomeres are in a different state of the contraction cycle. This is in agreement with the distinct thin filament organization observed at the M-line, where arrays of opposite polarity are in close apposition to each other without clear overlapping part (Fig. 3D).

## Conclusions

Our integrative approach permitted to directly visualize sarcomere organization in pristinely preserved cardiac muscle cells with molecular precision. Contracted sarcomeres, in which thin filaments are strongly bound to thick filaments, exhibit an overlap between thin filaments of opposite polarity in the M-line region, indicating that they slid past each other during contraction (fig. S10A). When the actomyosin binding is weaker, sarcomeres are found in an intermediate state, in which thin filaments of opposite polarity are in close apposition but do not overlap (fig. S10B). At the neonatal stage, contraction occurs despite an excess of thin filaments outside the trigonal positions of the thick-filament hexagonal lattice, including at the M-line where both actin polarities contribute to the overall packing (fig. S10A). Our work demonstrate that cryo-ET is able to reveal the polarity and functional states of the thin filaments *in situ*, and paves the way for structure determination of actomyosin interactions within cells.

## ACKNOWLEDGEMENTS

The authors thank W. Wan, F. Beck, P.S. Erdmann and V. Lucic for technical and computational assistance; R. Poincloux for critical reading of the manuscript and helpful comments. This work was supported by the Max Planck Society and by the German Center for Cardiovascular Research (DZHK)-Munich Partner Site (to R.T.B.). This preprint was formatted using the HenriquesLab bioRxiv template.

## AUTHOR CONTRIBUTIONS

L.B., J.S. and M.J. designed and interpreted experiments. S.S. prepared the cells and performed immunofluorescence imaging. L.B. performed cryo-FIB milling, cryo-ET, tomogram reconstruction, segmentation, and filament packing analysis with help from M.J. J.S. developed and applied the subtomogram averaging and polarity assessment approaches. R.T.B., W.B., P.S. and J.M.P. provided financial support and access to instrumentation. M.J. supervised the work and wrote the paper with contributions from all authors.

## DATA AND MATERIALS AVAILABILITY

The *in situ* structures of the cardiac thin filament have been deposited in the EM-DataBank under accession codes EMD-XXXX and EMD-YYYY.

## Materials and Methods

### Isolation of neonatal rat ventricular cardiomyocytes

Three-days-old (P3) wild-type Wistar rats were killed by decapitation and the ribcage was opened. The heart was removed by pulling with forceps and transferred into a 35-mm Petri dish containing phosphate-buffered saline (PBS). The remaining blood was pumped out of the heart and the great vessels. Both atria and the connective tissue were removed. For each preparation, the ventricles of four hearts were mixed together, cut into small pieces and digested using the Neonatal Heart Dissociation kit (Miltenyi Biotec, Bergisch Gladbach, Germany). Briefly, the enzyme mix 1 was pre-heated for 5 min at 37°C and added to the enzyme mix 2. The harvested and chopped ventricular tissue was transferred to the tube containing the enzyme mix. The suspension was gently pipetted every 10 min for 1h. Following enzymatic digestion, the cells were passed through a 70-μm cell strainer and centrifuged at 300g for 15 min. The pellet was resuspended in 10 ml of cardiomyocyte medium. This medium consists of a 1:1 mixture of Dulbecco’s modified Eagle medium (DMEM) and Ham’s F12 medium (Life technologies, #31330-038) containing 5% horse serum, 5% fetal calf serum (FCS), 20 μM Cytarabine (Ara-C) (Sigma Aldrich, #C1768), 3 mM sodium pyruvate (Sigma #S8636), 2 mM L-glutamine (Thermo Fisher Scientific, #25030-081), 0.1 mM ascorbic acid (Sigma #A4034), 1:200 insulin-transferrin-selenium-sodium pyruvate (Invitrogen, #51300044), 0.2 % bovine serum albumin (BSA) (Sigma Aldrich, #A7409) and 100 U/ml penicillin-streptomycin (Thermo Fisher Scientific, #15140122). The resuspended pellet was pre-plated onto a non-coated cell culture dish for 90 min to reduce the number of cardiac fibroblasts. The supernatant was transferred to a 15 ml tube and centrifuged at 300g for 15 min. The pellet containing the neonatal rat ventricular cardiomyocytes from four hearts was resuspended in 4 ml of cardiomyocyte medium (that is, 1 heart eq./ml). Moreover, primary adult mouse ventricular cardiomyocytes were isolated following the protocol from Ackers-Johnson et al. (34).

### Cardiomyocyte culture on EM grids and vitrification

Gold EM grids with Quantifoil R 2/1 holey carbon film (Quantifoil Micro Tools GmbH, Grossloebichau, Germany) were glow-discharged, sterilized by UV irradiation for 30 min and coated with fibronectin (10 μg/ ml, Merck, #341631). Six to eight EM grids were placed in 35-mm Petri dishes and 300 μl of cell suspension was added (that is, ~1/3 heart eq. per dish). The cells were cultured at 37°C in 5% CO_2_ for two days. To verify that the cells exhibit spontaneous rhythmic contractions prior to cryo-fixation, the grids were imaged at room temperature (RT) by bright field using a 20x (air, NA 0.8) Plan Achromat objective (Carl Zeiss, Jena, Germany) of a CorrSight microscope (Thermo Fisher Scientific). The grids were plunge-frozen using a Vitrobot Mark IV (Thermo Fisher Scientific) with 80% humidity and 10s blotting time in a 2:1 ethane/propane mixture cooled by liquid nitrogen.

### Immunofluorescence microscopy

Cells used for immunofluorescence microscopy were cultured on a chambered coverslip with 8 wells (IBIDI #80826, Gräfelfing, Germany) coated with fibronectin (10 μg/ml at 37°C for 1h). After two days, the cells were rinsed in PBS, placed on ice and fixed in 4% paraformaldehyde (PFA) for 15 min. After PBS wash, the cells were permeabilized with 0.1% Triton X-100/PBS for 10 min and blocked with 3% BSA/PBS at 4°C for at least one 1h. Subsequently, the cells were incubated at 4°C with one of the following mouse monoclonal primary antibodies: anti-heavy chain cardiac myosin (Abcam, #ab50967), anti-α-actinin (Sigma-Aldrich, #A7811) or anti-cTnI (Thermo Fisher Scientific, #MA5-12960). The next day, the cells were washed with PBS (3 × 5 min) and incubated with an anti-mouse secondary antibody (Alexa Fluor™ 488 goat IgG (H+L), Thermo Fisher Scientific, #A11029) at RT for 1h in the dark. The cells were washed with PBS (2 × 5 min), stained with DAPI (1:6,000 in PBS) at RT for 15 min in the dark, washed again in PBS (2 × 5 min) and stored at 4°C. Confocal imaging was performed with a Zeiss LSM 780 confocal laser scanning microscope (Carl Zeiss, Oberkochen, Germany) equipped with a 40x (oil, NA 1.4) Plan-Apo objective (Carl Zeiss, Jena, Germany). The resulting images were analyzed with ImageJ software version 1.52p (https://imagej.net/Fiji).

### Cryo-FIB milling

Plunge-frozen grids were clipped into Autogrids modified for cryo-FIB milling (35, 36). The Autogrids were mounted into a custom-built FIB shuttle cooled by liquid nitrogen and transferred using a cryo-transfer system (PP3000T, Quorum Technologies) to the cryo-stage of a dual-beam Quanta 3D FIB/SEM (Thermo Fisher Scientific) operated at liquid nitrogen temperature. During the loading step, the grids were sputter-coated with platinum in the Quorum prep-chamber (10 mA, 30s) to improve sample conductivity. To reduce curtaining artifacts, they were subsequently sputter-coated with organometallic platinum using the gas injection system (GIS, Thermo Fisher Scientific) in the microscope chamber operated for 8s at 26°C. Lamellas were prepared using a Gallium ion beam at 30 kV and stage tilts of 18-20°. 8-12 μm wide lamellas were milled in a step-wise manner using high currents of 0.5 nA for rough milling that were gradually reduced to 30 pA for fine milling and final cleaning steps. The milling process was monitored using the electron beam at 5 kV and 11.8 pA or 3 kV and 8.87 pA. For Volta phase plate (VPP) imaging, the lamellas were additionally sputter-coated with a platinum layer in the Quorum prep-chamber (10 mA, 3s).

### Cryo-ET

The lamellas were loaded in a Titan Krios transmission electron microscope (Thermo Fisher Scientific) equipped with a 300-kV field-emission gun, VPPs (Thermo Fisher Scientific), a post-column energy filter (Gatan) and a 4k × 4k K2 Summit direct electron detector (Gatan) operated using SerialEM (37). The VPPs were aligned and used as described previously (38). Low-magnification images were captured at 6,500x. High-magnification tilt series were recorded in counting mode at 42,000x (calibrated pixel size of 0.342 nm), typically from −50° to 70° with 2° steps, starting from a pre-tilt of 10° to correct for the lamella geometry, and a total dose of 100-120 e^−^/Å^2^. Three of the tilt series were collected unidirectionally with the VPPs at a target defocus of 0.5 μm. Ten of the tilt series were recorded without the VPP using a dose-symmetric tilt-scheme (39) and a target defocus range of −3.25 to −5 μm.

### Tomogram reconstruction

Frames were aligned using MotionCorr2 (40) and dose-filtered by cumulative dose using the exposure-dependent attenuation function and critical exposure constants described in (41) and adapted for tilt series in (42, 43). Tilt series were aligned using patch tracking in IMOD (44) and tomograms were reconstructed with a binning factor of 4 (13.68 Å per pixel) using weighted back-projection. For visualization, tomograms were filtered using a deconvolution filter (https://github.com/dtegunov/tom_deconv). For subtomogram averaging, contrast transfer function (CTF) parameters for each tilt were estimated by CTFFIND4 (45). 3D-CTF corrected tomograms were generated using NovaCTF (46) and binned by a factor of 2 (6.84 Å per pixel).

### Membrane and filament segmentation

The 4 × binned tomograms (pixel size of 13.68 Å) were used for segmentation. Membranes were generated automatically using TomoSegMemTV (47) and refined manually in Amira (Thermo Fisher Scientific). Thin and thick filaments were traced automatically in Amira using an automated segmentation algorithm based on a generic cylinder as a template (48). The cylindrical templates were generated with a length of 42 nm and diameters of 8 and 18 nm for the thin and thick filaments, respectively.

### Nearest neighbor analysis and local packing analysis

The analysis developed in (25) was adapted to describe quantitatively the 3D organization of the cardiac myofilaments. A total of 4,770 thin filaments and 1,648 thick filaments extracted from 9 tomograms were used. Data analysis was performed in MATLAB (The MathWorks) using the coordinates of the filament centerlines exported from Amira as input. The coordinates of the filaments were resampled every 3 nm to give the same weight to every point along a filament.

#### Nearest neighbor analysis

The local direction at each point was evaluated as the local tangent of the filament at that point. For every point within a filament, the closest point of each neighboring filament was characterized by its distance, d, and relative orientation, ϑ, with respect to the reference point. The analysis was performed for filaments of the same type (that is, only thick filaments or only thin filaments) as well as for filaments of one type relative to the other (that is, neighboring thin filaments for the thick filaments, and reciprocally). The occurrences of (d, ϑ) were represented by a 2D histogram. The peak occurring at small ϑ (below 10°) indicates the presence of equidistant and nearly parallel filaments. The mean interfilament spacing was obtained from the distances with a number of occurrences higher than two third of the peak maximum (fig. S4).

#### Local packing analysis

A local reference frame (e1, e2, e3) was defined at every point along a filament as follows: e2 points in the local direction of the filament, e1 points in the direction of the nearest neighbor, and e3 is the cross-product of e1 and e2 (see (25)). (X, Y, Z) are the coordinates of a neighbor position in (e1, e2, e3). The local reference frames (e1, e2, e3) of all points along the filaments were aligned, and the occurrences of (X, Z) in the e1e3 plane within the distance range for parallel filaments were represented by a 2D histogram (Fig. 2, top).

### Subtomogram averaging and polarity assessment

#### Subtomogram sampling

Ten tomograms, acquired with a dose-symmetric tilt-scheme and a target defocus range of −3.25 to −5 μm, were used for subtomogram averaging and polarity assessment. The coordinates of the thin filaments extracted from Amira were resampled every 1.38 nm (corresponding to half of the axial rise per actin subunit in the actin filament) along the filament centerline using scripts described in (25). This was instrumental to correctly allocate every actin subunit during subtomogram alignment (42). Principal component analysis was used to generate a common direction for all the filaments in the network upon resampling (that is, filament coordinates were reordered so that the dot product between the direction of each filament and the first principal component was positive). The local tangent at each point along a filament served to calculate initial Euler angles (φ,ϑ,ψ) in the zxz-convention using scripts from (25) and TOM Toolbox (49). As a result, the filament centerline was aligned with the z-axis, allowing the φ angle to describe the in-plane rotation. Since this rotation cannot be determined directly from the tomogram, φ was randomized every 30°. Furthermore, subvolumes belonging to the same filament were assigned a unique identifier, which was used during the assessment of filament polarity exclusively.

#### De novo reference

Subtomogram averaging was performed using STOPGAP (https://github.com/williamnwan/STOPGAP). An initial reference was generated *de novo* from a single tomogram as follows. 84 thin filaments located on one side of a Z-disk and belonging to the same bundle (shown in orange in fig. S6A) were selected, based on the assumption that they had the same polarity. 20,259 subtomograms were extracted from the resampled positions using a binning factor of 2 (6.84 Å per pixel) and a box size of 128³ pixels³ and a starting reference was generated by averaging all subvolumes. Due to the in-plane randomization, the initial structure resembles a featureless cylinder (fig. S6B). Several rounds of global alignment of the in-plane angle followed by local refinement of all Euler angles were conducted. Shift refinements were limited to 1.38 nm in each direction along the filament centerline. This averaging approach resulted in the emergence of the helical pattern of the thin filament (fig. S6B). Unbinned subvolumes were extracted using a box size of 128³ pixels³ and aligned iteratively, resulting in a refined structure of F-actin in complex with Tpm at a resolution of 20.7 Å (fig. S6, B and C). At this resolution, the positions of the actin subunits with respect to the two Tpm strands allowed to distinguish between F-actin structures of opposite polarities (fig. S6D). Comparison with a pseudoatomic model of the F-actin-Tpm complex (10) permitted to allocate the barbed (+) and pointed (−) end of the *de novo* F-actin structure (fig. S6E).

#### Polarity assessment

The 20.7 Å structure of F-actin in complex with Tpm was rotated by 180° around the x-axis to generate a second structure with opposite polarity (Fig. 3A and fig. S6D). Both structures were used to assess the polarity of the thin filaments in each tomogram using multireference alignment (fig. S7A). Each subvolume within a tomogram was extracted with a binning factor of 2 (6.84 Å per pixel) using a box size of 64³ pixels³ and aligned against both references independently. The scores of all the subtomograms belonging to the same filament were compared using unpaired Student’s t-test (fig. S7B). On a total of 4,438 filaments (corresponding to 594,317 subvolumes) from 10 tomograms, 69% showed significant differences between the two directions (p < 0.05) and were assigned the respective polarity with confidence (fig. S7C). These filaments were kept for further processing. Polarity assignment was visualized with arrows pointing in the direction of the barbed ends of F-actin using the Place Object tool in UCSF Chimera (50, 51).

#### Subtomogram averaging of the thin filament

To generate a dataset with uniform polarity, the initial Euler angles from the subvolumes of opposite polarity were rotated by 180° around the x-axis. 409,896 subtomograms were extracted with a binning factor of 2 (6.84 Å per pixel) using a box size of 64³ pixels³. The 20.7 Å *de novo* structure of F-actin in complex with Tpm (fig. S6B) was low-pass filtered and used as initial reference. Global alignment of the in-plane angle was followed by several rounds of successive local refinement of the in-plane angle and all Euler angles. Simultaneously, shifts were refined within ± 1.38 nm along the filament centerline. Once all the positions were refined, distance thresholding was used to remove oversampled positions, keeping only one subtomogram per actin subunit. 160,617 unbinned subvolumes were extracted using a box size of 128³ pixels³. The dataset was split into two half sets that were refined independently.

To uncover potential changes in the azimuthal position of the Tpm strands between sarcomeres, each tomogram was then processed separately using the initial reference without Tpm density. The data shown in Fig. 3D yielded a subtomogram average in which the Tpm density was significantly shifted on F-actin. The other tomograms provided a comparable structure and the corresponding 133,587 particles were merged and further refined. The final maps were sharpened, weighted by the FSC and filtered to their respective resolution (FSC_0.143_ = 17.3 and 13.2 Å, respectively, fig. S9A). Local resolution calculations were performed using RELION 3.0 (52) (fig. S9C). All maps were visualized using UCSF Chimera or UCSF Chimera X (53). Analysis of the distances and in-plane rotation between refined subvolumes belonging to the same filament permitted to determine the mean rise and angular twist of the F-actin structure.

Several pseudoatomic models containing F-actin in complex with Tpm in distinct functional states (10, 18) were docked into the maps using rigid body-fitting in UCSF Chimera. The fit was limited to the actin region to reveal the functional state matching the Tpm density best.

---

**Fig. S1.**
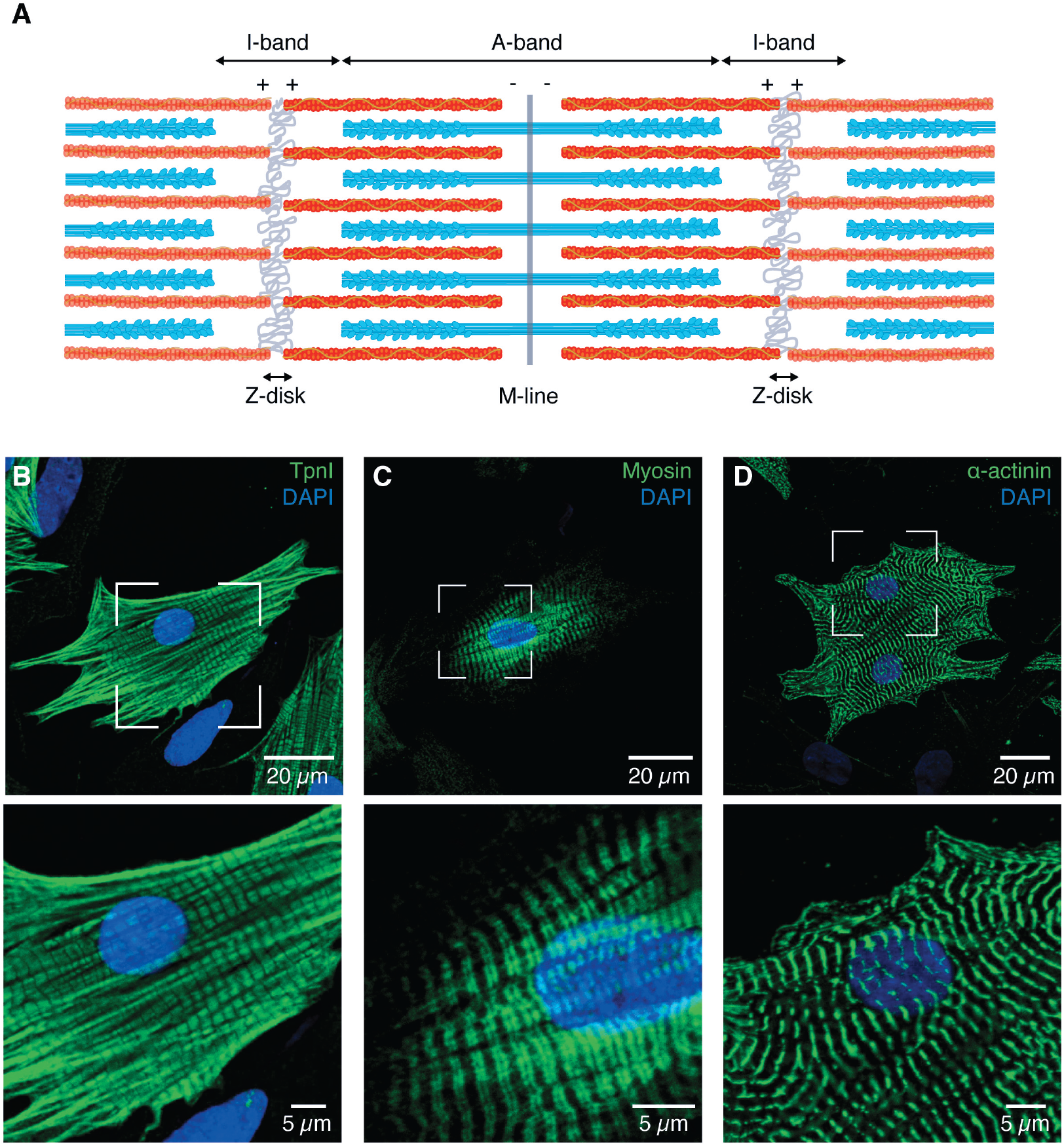
Sarcomere organization at the microscale. (A) Schematic representation of adjoining micrometer-sized sarcomeres within a myofibril, showing the organization of the thick (cyan) and thin (red) filaments in this basic contractile unit of striated muscle. (B-D) Thin (B) and thick (C) filaments of neonatal rat cardiomyocytes assemble into myofibrils, which align with the main axes of the star-like shaped cells and display regularly spaced Z-disks (D). TpnI, troponin I.

**Fig. S2.**
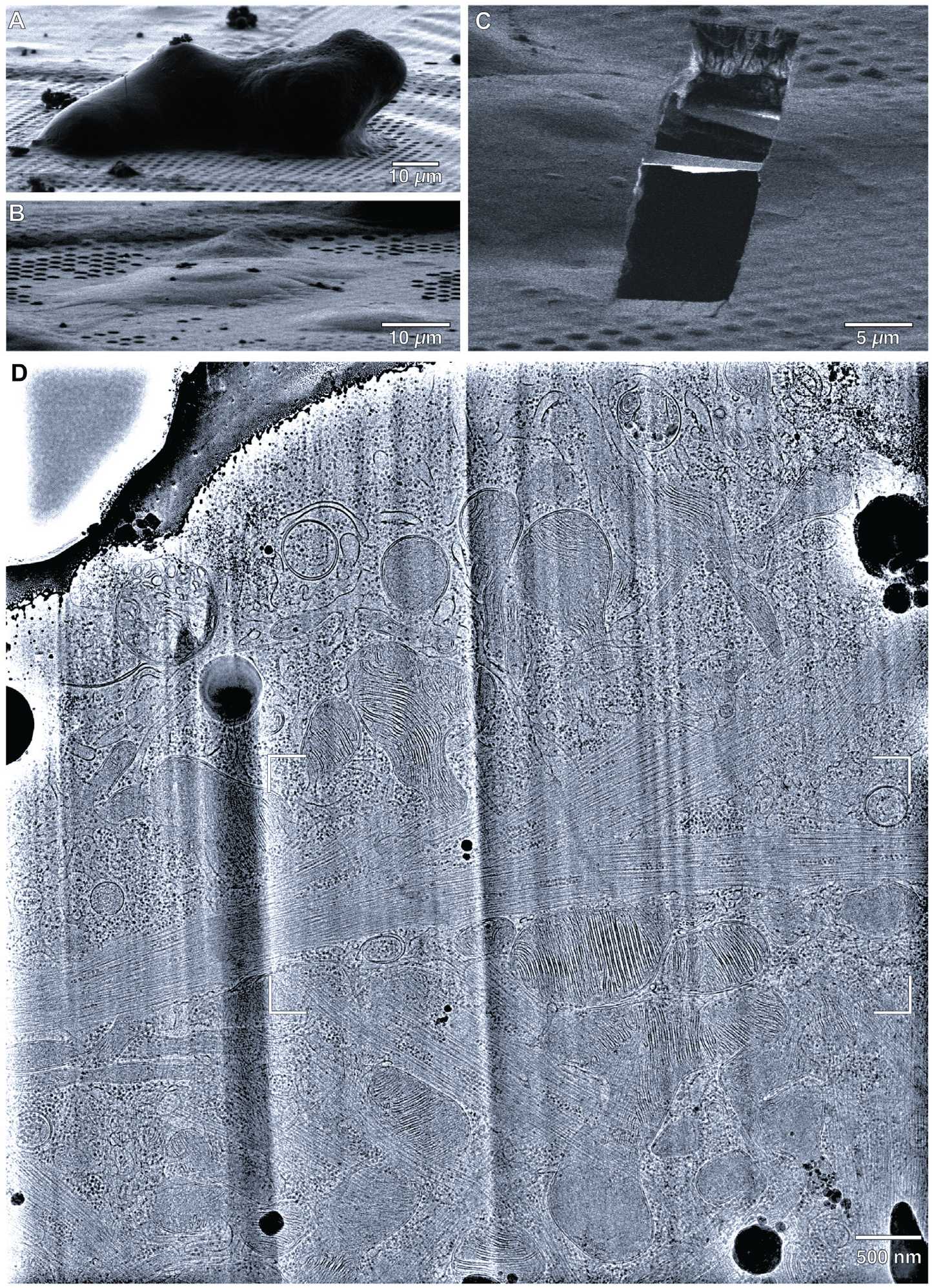
Cryo-focused ion beam sample preparation of neonatal cardiomyocytes for cryo-ET imaging. (A) Scanning electron microscope (SEM) image of a frozen-hydrated adult mouse cardiomyocyte on an EM grid. Vitrification by plunge-freezing is not suitable for these cells because their thickness is larger than 10 μm. (B) Neonatal rat cardiomyocytes are thin, allowing optimal vitrification by plunge-freezing without the need for cryoprotectant. (C) 200-nm-thick lamella prepared in a plunge-frozen neonatal cardiomyocyte and imaged by FIB-induced secondary electrons. (D) Full TEM image of a neonatal cardiomyocyte lamella containing the cropped part shown in Fig. 1C.

**Fig. S3.**
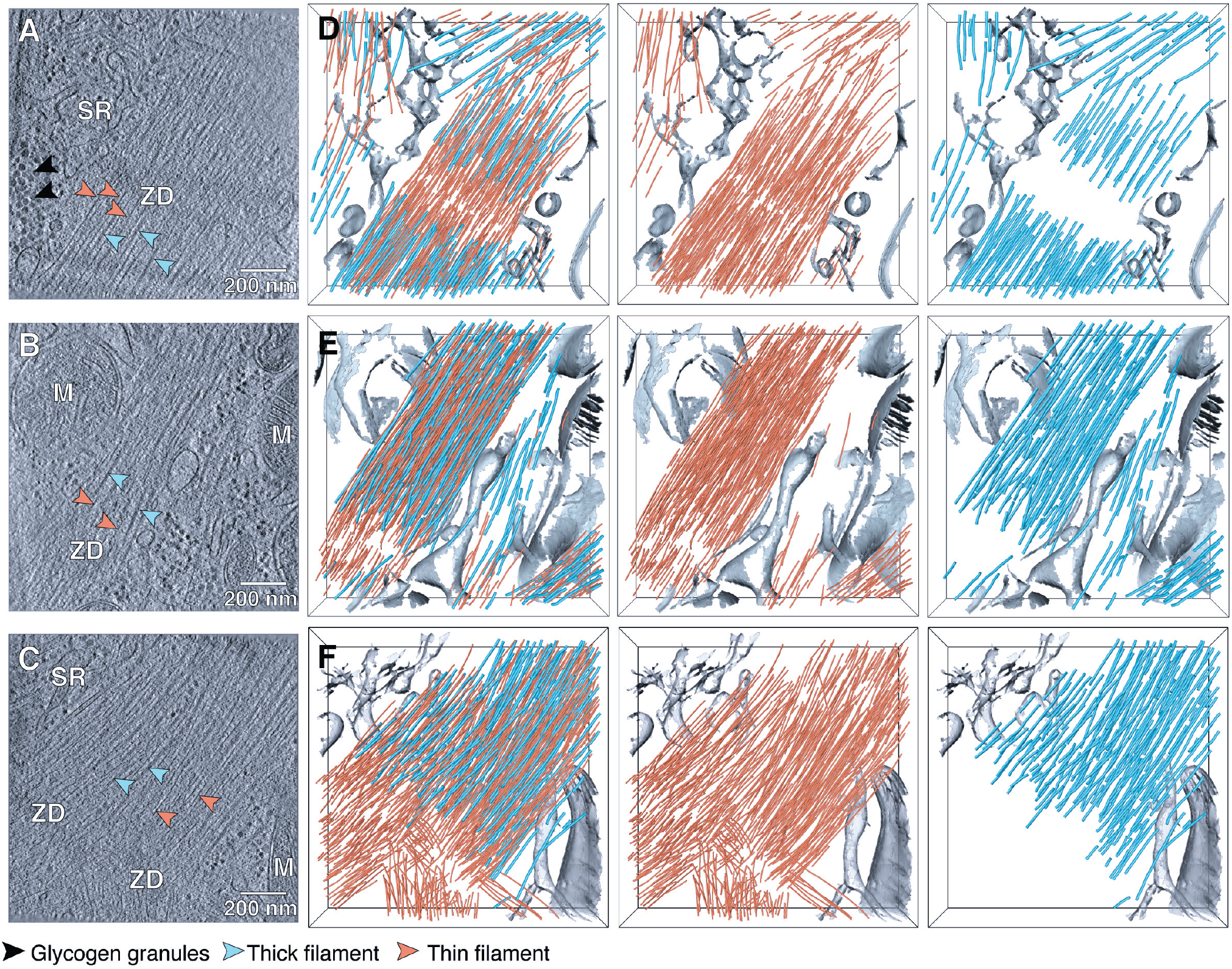
Nanoscale sarcomere organization revealed by *in situ* cryo-ET. (A-C) Slices from tomographic volumes acquired in frozen-hydrated neonatal rat cardiomyocytes, revealing the myofibrillar environment. ZD, Z-disk; M, mitochondrium; SR: sarcoplasmic reticulum. (D-F) Corresponding 3D segmentation of the cellular volumes, showing the nanoscale organization of the thin (orange) and thick (cyan) filaments within unperturbed sarcomeres.

**Fig. S4.**
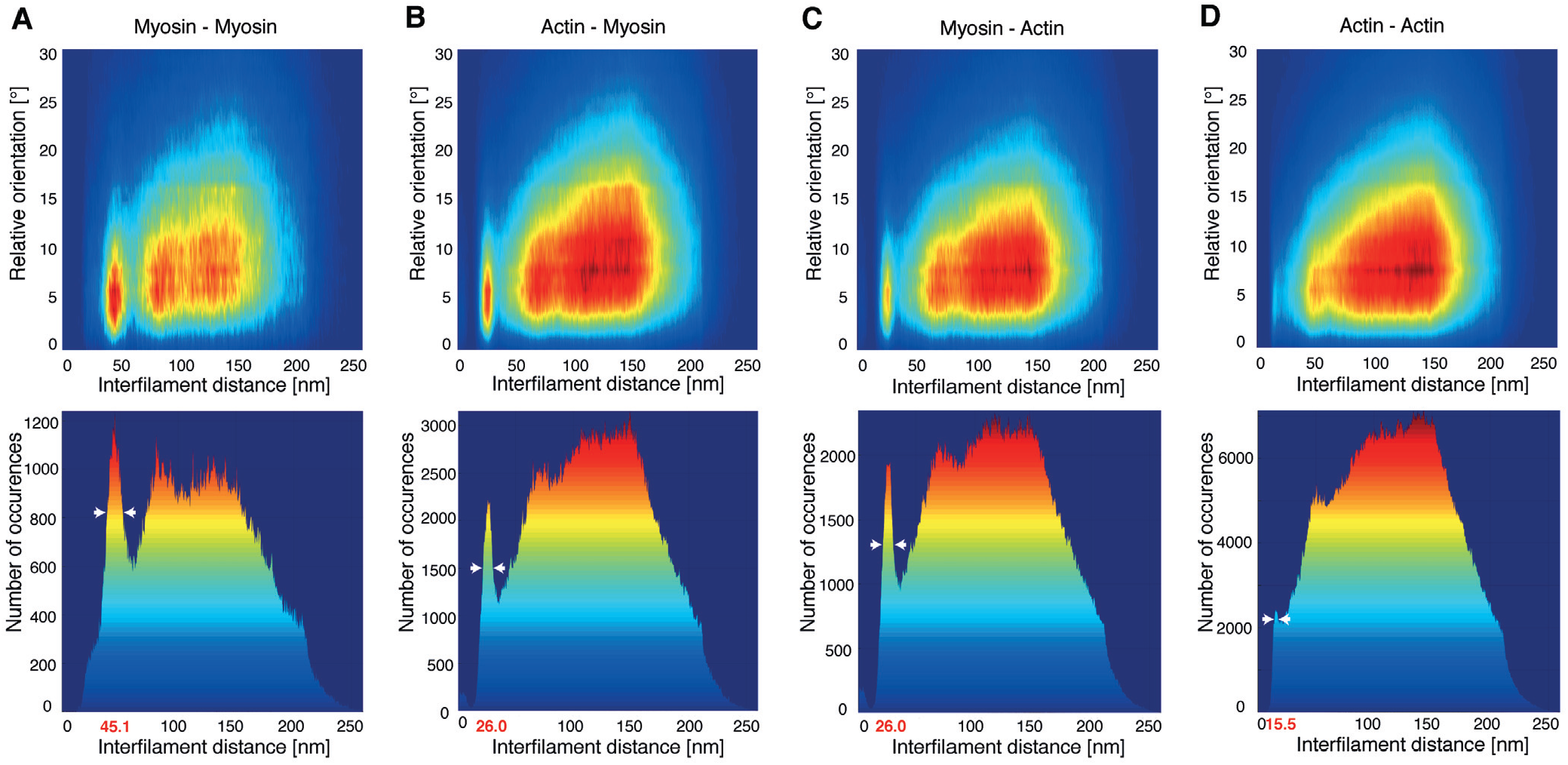
Nearest-neighbor analysis of myofilaments in neonatal rat cardiomyocytes. (A-D) Top: 2D histogram of interfilament distances and relative orientations between thick filaments (“Myosin”; A), between thick and thin filaments (“Actin”; B and C), and between thin filaments (D). Bottom: corresponding histograms showing the number of occurrences of the interfilament distances. Thick filaments have an interfilament spacing of ~ 45 nm (A) and are found at ~ 26 nm from the thin filaments (B and C). Thin filaments have their nearest actin neighbors at ~ 15.5 nm (D). White arrow heads indicate the distance range for parallel filaments.

**Fig. S5.**
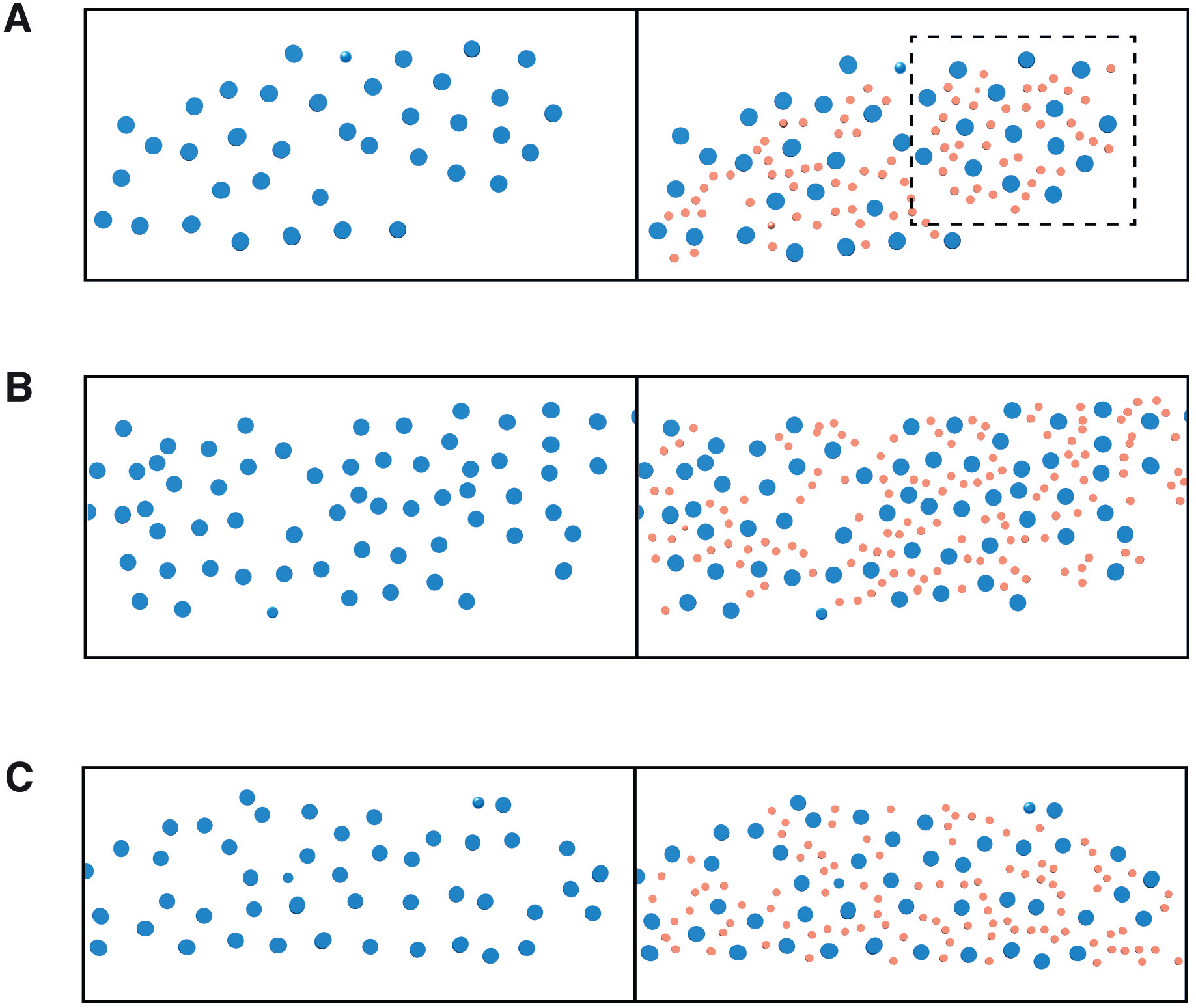
Examples of cross-sections through myofibrils of neonatal rat cardiomyocytes. (A-C) Thick filaments (blue) are hexagonally packed and surrounded by thin filaments (orange, right) at the trigonal positions of the lattice and outside of them. The framed area in (A) is shown in Fig. 2C, bottom. The slices were obtained from the myofibrils shown in fig. S3E (A), Fig. 1E (B) and fig. S3F (C).

**Fig. S6.**
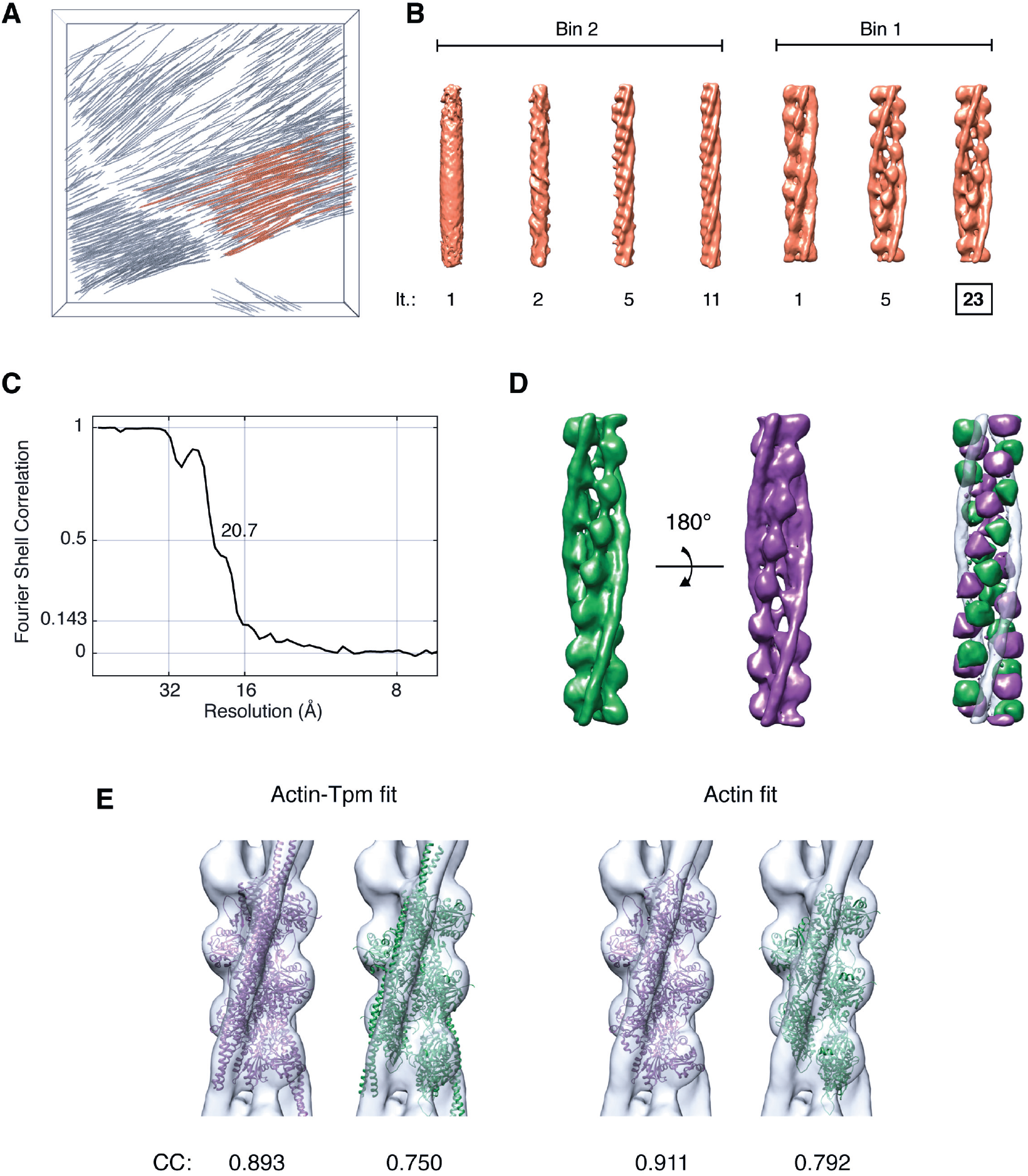
*De novo* structure generation from cryo-ET data of cardiac thin filaments. (A) Thin filaments (orange) from the tomogram shown in Fig. 1D and movie S2 used for the generation of a *de novo* reference. (B) Iterative alignment using 2 × binned (“Bin2”) and unbinned (“Bin1”) data resulted in the emergence of the helical pattern of the thin filament and yielded a structure of F-actin in complex with Tpm resolved at 20.7 Å. It: Iteration number. (C) Corresponding Fourier shell correlation plot. Since the data was not processed using gold-standard refinement, a cutoff of 0.5 was used. (D) Difference map (right) generated by aligning the Tpm densities of the filament structures in opposite orientations (left; green and purple, respectively), revealing the actin positions associated to each polarity. (E) Both orientations (same color code as in (D)) of the pseudoatomic model of the F-actin-Tpm complex (pdb 5jlh) docked into the *in situ* map (grey) shown in purple in (D) (left), and corresponding fits using the actin densities only (right). The cross-correlation (“CC”) results permitted to allocate the barbed (+) and pointed (−) end of the *de novo* F-actin structure (Fig. 3A).

**Fig. S7.**
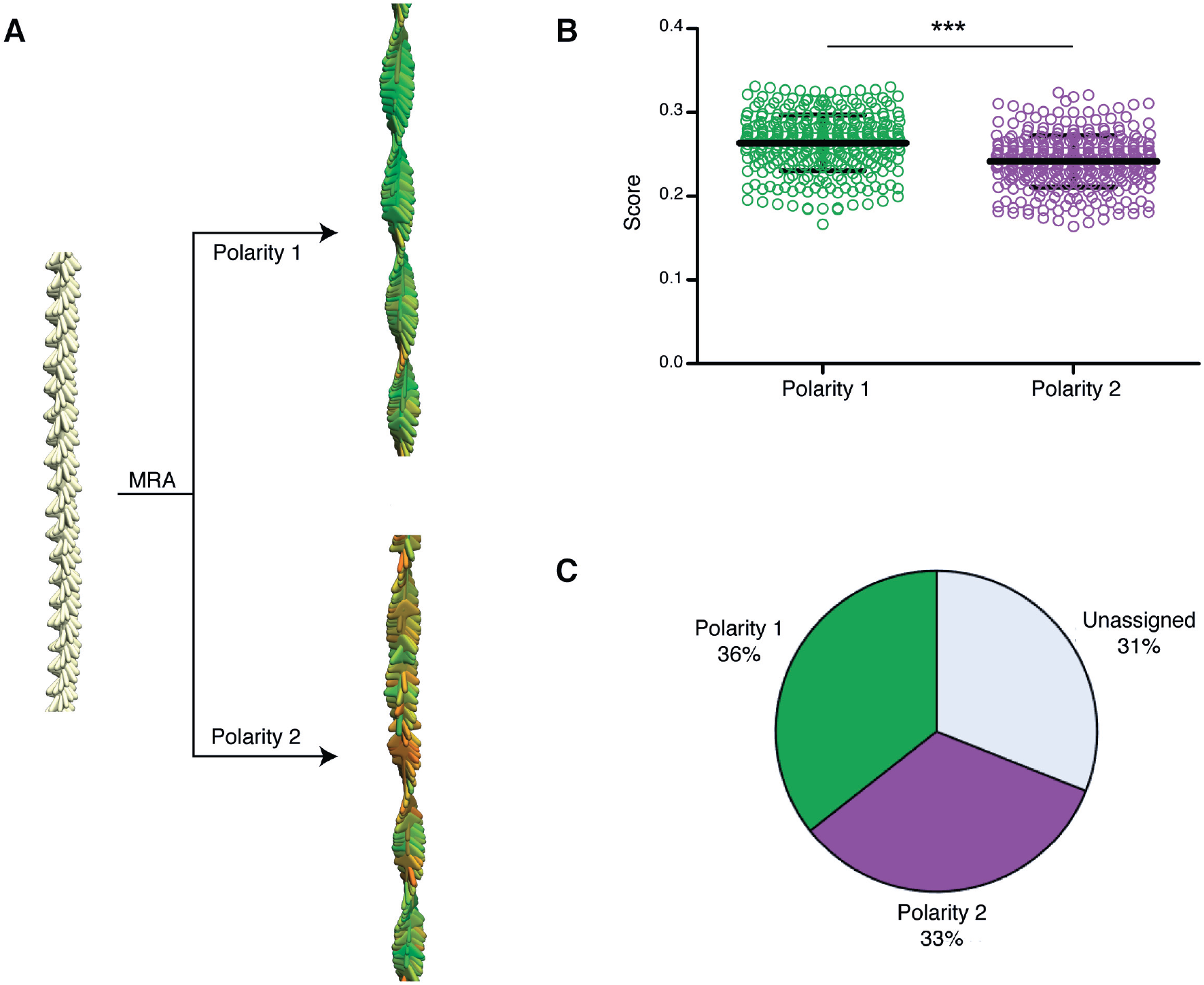
Workflow for the polarity assessment of the thin filaments imaged by cryo-ET. (A) Multireference alignment (“MRA”) procedure applied to one filament with oversampled positions aligned along the z-axis and in-plane randomization represented by yellow arrows (left). Each position is aligned independently against a reference of each polarity (in green and purple, respectively), resulting in refined orientations and scores (high scores, green; low scores, red) (right). Refinement against the green structure (“Polarity 1”) leads to uniform helical pattern, whereas alignment against the purple structure (“Polarity 2”) yields irregularities in the helical pattern. (B) Corresponding scores obtained for the single filament under each polarity, with the mean and standard deviation indicated. Unpaired Student’s t-test revealed significant differences between both polarities (***: p < 0.001). (C) Pie chart representing the results of the polarity assignment for the entire dataset of 4,438 thin filaments with p < 0.05.

**Fig. S8.**
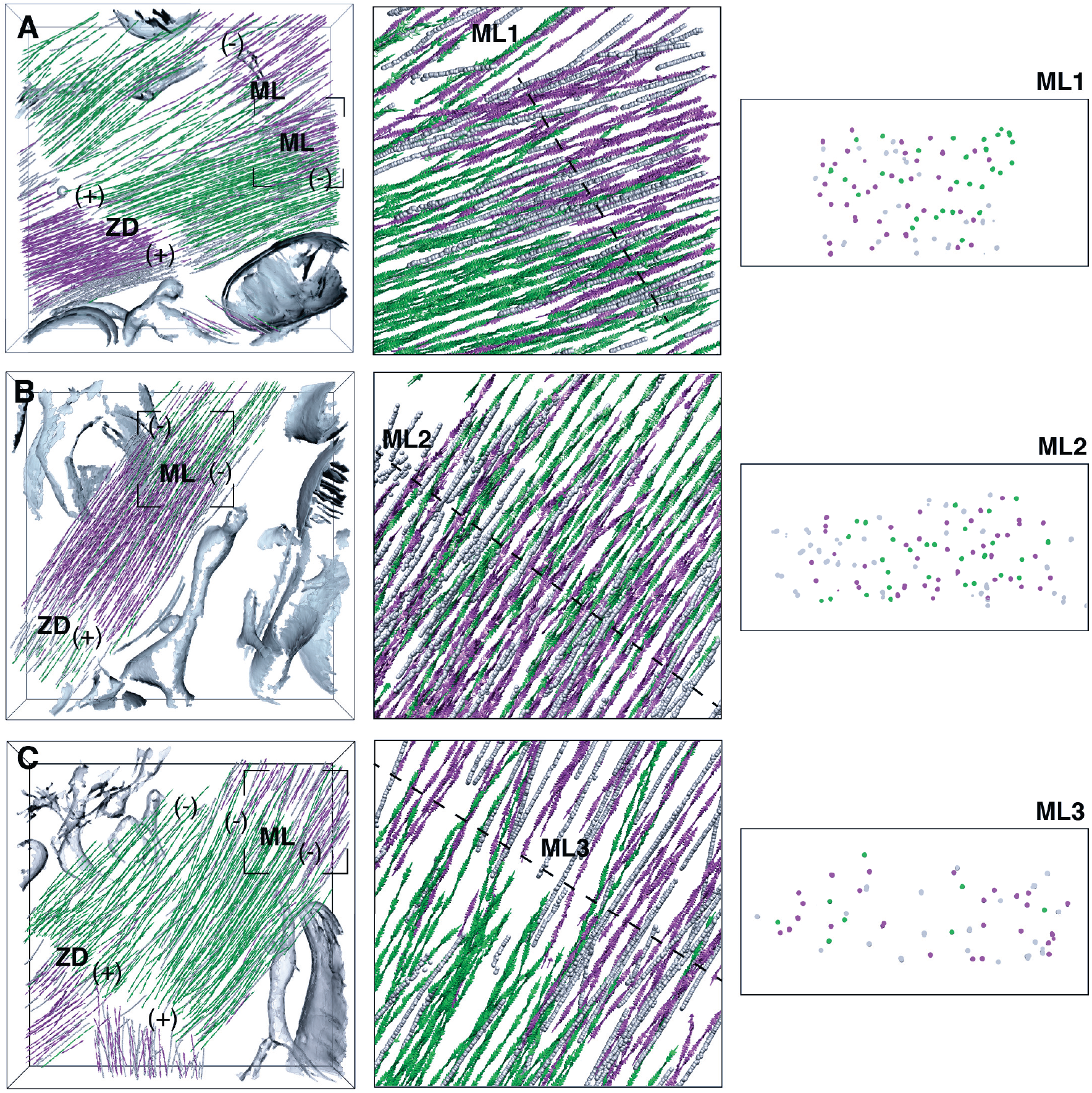
Polarity assignment and packing of cardiac thin filaments at the center of the sarcomere. (A-C) Same examples as in Fig. 3 showing the unassigned filaments in grey, which correspond to 23% of the filaments in these datasets. Thin filaments with assigned polarity are represented by colored arrows pointing toward the barbed (+) end. ZD, Z-disk; ML, M-line. Insets: Thin filament orientations in the framed M-lines (middle) with their respective cross-sections (right).

**Fig. S9.**
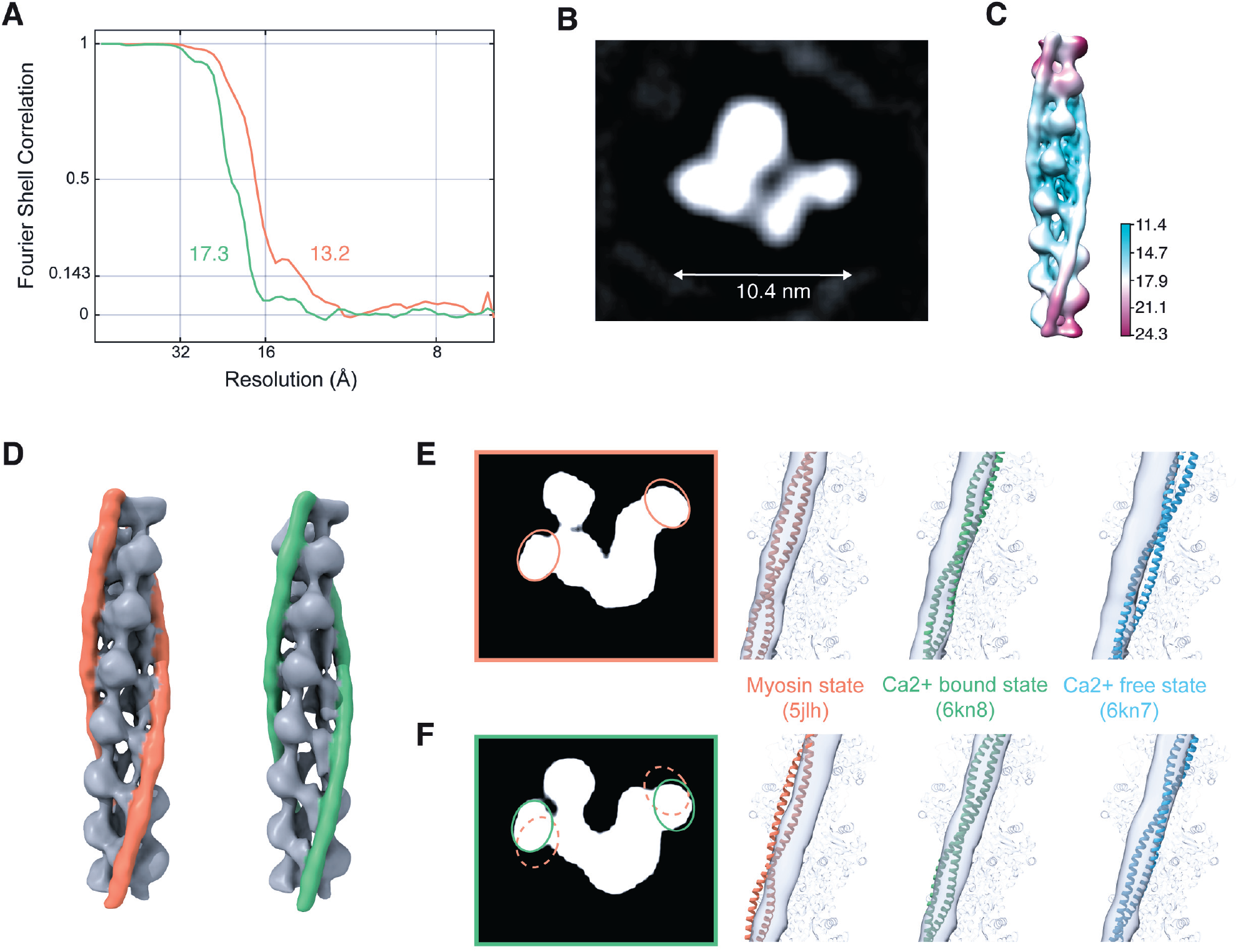
Resolution and functional state of the native cardiac thin filament structures. (A) Gold-standard FSC plots of the subtomogram averages shown in Fig. 4A (orange curve) and E, middle (green curve). (B) x-y slice through the 13.2 Å map. (C) Local resolution estimation. Due to the averaging procedure, the resolution of the filament is the highest in the central subunits. (D-F) x-y slices through the 13.2 Å (E) and 17.3 Å (F) resolution structures shown in (D), in which the Tpm densities are highlighted in orange and green, respectively, and close up into the fits of the pseudoatomic models in different states into the maps.

**Fig. S10.**
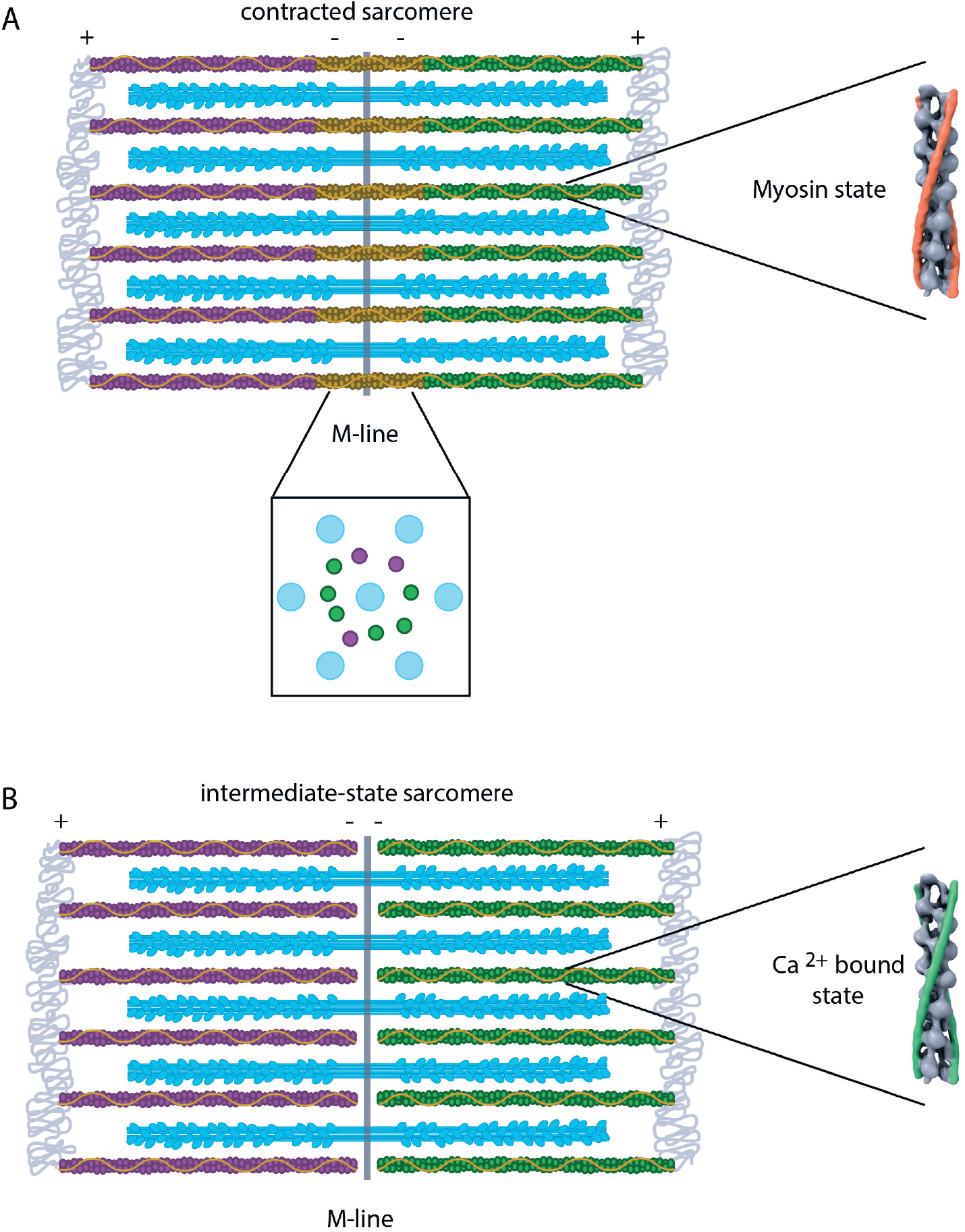
Thin filament organization during sarcomere contraction. (A) Contracted sarcomeres, in which thin filaments are strongly bound to thick filaments, exhibit an overlap between thin filaments of opposite polarity in the bare zone region. (B) When the actomyosin interaction is weaker, sarcomeres are found in an intermediate state, in which thin filaments of opposite polarity are in close apposition but do not overlap.

